# Demonstration of *de novo* chemotaxis in *E. coli* using a real-time, quantitative, and digital-like approach

**DOI:** 10.1101/114207

**Authors:** Tzila Davidov, Naor Granik, Sharbel Zahran, Inbal Adir, Ofek Elul, Tal Fried, Asif Gil, Bar Mayo, Shilo Ohayon, Shiran Sarig, Nofar Shasha, Shirane Tsedef, Shani Weiner, Michal Brunwasser-Meirom, Alexandra Ereskovsky, Noa Katz, Beate Kaufmann, Yuri Haimov, Heidi Leonard, Ester Segal, Roee Amit

## Abstract

Chemotaxis is the movement of an organism in response to an external chemical stimulus. This system enables bacteria to sense their immediate environment and adapt to changes in its chemical composition. Bacterial chemotaxis is mediated by chemoreceptors, membrane proteins that bind an effector and transduce the signal to the downstream proteins. From a synthetic biology perspective, the natural chemotactic repertoire is of little use since bacterial chemoreceptors have evolved to sense specific ligands that either benefit or harm the cell. Here we demonstrate that using a combined computational design approach together with a quantitative, real-time, and digital detection approach, we can rapidly design, manufacture, and characterize a synthetic chemoreceptor in *E. coli* for histamine (a ligand for which there are no known chemoreceptors). First, we employed a computational protocol that uses the Rosetta bioinformatics software together with high threshold filters to design mutational variants to the native Tar ligand binding domain that target histamine. Second, we tested different ligand-chemoreceptors pairs with a novel chemotaxis assay, based on optical reflectance interferometry of porous silicon (PSi) optical transducers, enabling label-free quantification of chemotaxis by monitoring real-time changes in the optical readout (expressed as the effective optical thickness, EOT). We found that different ligands can be characterized by an individual set of fingerprints in our assay. Namely, a binary, digital-like response in EOT change (i.e. positive or negative) that differentiates between attractants and repellants, the amplitude of change of EOT response, and the rate by which steady state in EOT change is reached. Using this assay, we were able to positively identify and characterize a single mutational chemoreceptor variant for histamine that mediated chemotaxis comparably to the natural Tar-aspartate system. Our results demonstrate the possibility of not only expanding the natural chemotaxis repertoire, but also provide a new quantitative assay by which to characterize the efficacy of the chemotactic response.

## Introduction

Bacterial chemotaxis is the movement of bacteria in response to an external chemical stimulus. This system enables bacteria to sense their immediate environment and quickly adapt to changes in its chemical composition, moving away from repellents or towards attractants, thus assuring their survival. Furthermore, it is characterized by specific responses, high sensitivity, and a dynamic range (1, 2). Chemotaxis is facilitated by methyl-accepting chemotaxis protein (MCP or the chemoreceptor), which is typically a transmembrane protein that binds the effector through the periplasmic ligand binding site and transduces the chemotactic signal forward to downstream Che proteins (1, 3, 4). Most MCPs contain a poorly-conserved ligand-binding domain, one or more HAMP domains, and a conserved C-terminal cytoplasmic domain that contains the methylated glutamate residues. The level of methylation is controlled by the interplay between the methylesterase (methyl-removing) phosphorylated CheB and the methyltransferase CheR (1, 4). Binding a ligand causes a conformational change in the MCP receptor, which translates down the hairpin structure. An attractant facilitates increased methylation of the C-terminus domain, while a repellant causes the opposite effect to occur. These methylation states, in turn, lead the bacteria to either travel up a gradient for an attractant, or away from it with a repellant. The motion is controlled via a signal propagation pathway that originate with the signal transducer (CheW) and histidine kinase component (CheA), which can bind the tip of the conserved C-terminus domain of the different MCPs encoded within the bacterium. Together, the molecular components of chemotaxis can be thought of as a broad-band chemical detection apparatus for gradients of varying analytes (1, 3).

Bacterial chemotaxis is characterized by simplicity of design, plasticity, and broad range of chemical detection repertoire that is highly conserved across the prokaryotic kingdom (5, 6). In particular, *E. coli’s* chemotaxis system consists of only five chemoreceptors: Tsr (taxis to serine and repellents), Tar (taxis to aspartate and repellents), Trg (taxis to ribose and galactose), Tap (taxis to dipeptides) and Aer (taxis to oxygen) (1, 3). Due to the universality of the signal transduction pathway, this simple, yet optimized configuration suggests that chemotaxis can be readily manipulated by simply adding new chemoreceptors, which can communicate with the downstream signaling apparatus. One of the most intriguing characteristics of chemotaxis is that the bacteria do not detect a concentration of a particular chemical, but rather a gradient in chemical compositions. This characteristic implies that chemotaxis has the potential to be an alternative for detection of chemicals in solutions, as gradients can be manipulated independently of analyte concentrations. As a result, several attempts in recent years have been made to increase the chemotaxis repertoire of *E. coli. (7, 8)* In all cases, the studies focused on altering the chemoreceptors in such a way that might result in a change of the specificity of the LBD and cause it to sense novel chemo-effectors. These attempts, although share the same goal, were based on two fundamental approaches: Lin *et al*(8) showed that *E. coli’s* Tar receptor can be re-specified to respond to cysteic acid, phenyalanin, and N-methyl-asparate using error-prone PCR on the periplasmic ligand binding domain, and with follow up selection experiments. Alternatively, two studies(9, 10) have shown that it is possible to make functional chemoreceptor chimeras in *E. coli* using the ligand binding domains of *Pseudomonas aeruginosa* PctA, PctB, and PctC chemoreceptors and the native Tar chemoreceptor. Notably, Bi *et* al.(7) recently developed a more systematic approach to engineering chemoreceptor chimeras in *E. coli,* thus expanding the synthetic chemotactic repertoire of *E. coli* further.

Here, we show that a combined computational design and a real-time quantitative detection method can be used to further expand the chemotactic repertoire of *E. coli* beyond naturally occurring ligand-binding domains. To do this, we implemented a two-pronged approach: first, we computationally designed novel chemoreceptors using the “Rosetta” bioinformatics software suite, and second, we developed a novel chemotaxis detection tool, which facilitates rapid, quantitative, and “digital-like” detection of attractants and repellants by monitoring the light reflective interference from nanostructured PSi-based optical transducer. (11) We demonstrate the feasibility of our approach, by using Rosetta to produce several candidate Tar-LBD mutants that had a high probability of binding histamine. We then show using “digital chemotaxis” and more traditional chemotaxis assays that one of the variants that received a high score by Rosetta was indeed able to respond to histamine in a specific manner.

## Materials and Methods

### Materials

Aspartate, histamine, methionine, N’-(3-triimethoxysilylpropyl) diethylenetriamine (DETAS), succinic anhydride, N,N’-diisopropylcarbodiimide (DIC), N-hydroxysulfosuccinimide (NHS), lectin from *Triticum vulgaris* (termed as wheat germ agglutinin or WGA), and acetonitrile were purchased from Sigma-Aldrich, Israel. Absolute ethanol, potassium and phosphate salts, dimethyl sulfoxide (DMSO), and diisopropylethylamine (DIEA) were supplied by Merck, Germany. Acetic acid, NaCl, and MgSO_4_ were supplied by Bio-Lab Ltd, Israel. Bacto agar and tryptone were supplied by BD Biosciences, USA. Low melt SeaPlaque agarose was purchased from Lonza. EDTA was supplied by JT Baker. Sodium benzoate was supplied by the Fluka brand of Honeywell (USA). Bio-assay buffer (BA) was composed of 0.5 mg/mL tryptone, 0.1 M NaCl, 1 M MgSO_4_ in 0.01 M PBS supplemented with 0.03% glycerol. TB-based medium was comprised of 10 g/L tryptone and 10 g/L NaCl. Motility buffer was composed of 0.1 M K_2_HPO_4_, 0.1 M KH_2_PO_4_, 0.1 mM EDTA, and 0.001 mM methionine in double-distilled water (ddH_2_O, 18 MΩ). Phosphate buffered saline (PBS) pH 7.2 was prepared by dissolving 17 mM KH_2_PO_4_, 27 mM KCl, 52 mM Na2HPO_4_, and 1.4 M NaCl in ddH_2_O.

### Silicon diffraction grating preparation

Photonic chips containing Si diffraction gratings in the form of arrayed micron-sized square wells were fabricated by standard lithography and reactive ion etching techniques (Micro-Nano Fabrication Unit, Technion - Israel Institute of Technology) and mechanically diced into 10 mm by 5 mm chips by an automated dicing saw (Disco, Japan). Lectin from *tritium vulgaris,* commonly referred to as wheat germ agglutinin (WGA), was immobilized on the Si diffraction gratings as previously described.(12) Briefly, the chips were thermally oxidized in ambient air at 800 °C for 1 hour in a Lindberg/Blue Split-Hinge furnace (Thermo Scientific, USA), followed by silanization in 2% DETAS (50% ethanol in ddH_2_O, acidified with 0.6% acetic acid) for 1 h. Chips were washed with ethanol and dried under a stream of nitrogen. Carboxylation was performed by incubation in a solution comprised of 1% succinic anhydride (in acetonitrile with 4% DIEA) for 3 h followed by washing with ethanol and drying with nitrogen. Amine activation was promoted by incubation in 129 mM DIC and 87 mM NHS constituted in acetonitrile for 1 h. After rinsing with ethanol, chips were stored at 4 °C overnight in WGA solution (1 mg/mL, 10% DMSO) to promote lectin immobilization onto the Si diffraction grating.

### Bacterial strains and plasmids

Derivative of *E. coli* RP437 strain UU1250 (Δ*tsr-7028* Δ*(tar-tap)5201* Δ*trg-100* Δ*aer-1, kan^r^)* (13) was used as a parental strain to construct all the chemotactic strains in this work, and were obtained from Dr. Vaknin (Hebrew University). ΔZras and Δfilm B275 strains were used as a positive and negative control strains, respectively, and were obtained from Prof. Eisenbach (Weizmann Institute). PSB1C3 and PSB1A3 plasmids were obtained from (International Genetically Engineered Machine) iGEM 2016 distribution kit.

### PctA-Tar chimera construction

To construct the chimeric chemoreceptor, the LBD and HAMP domains of the *E. Coli* Tar were replaced by the respective domains of the PctA receptor, as was previously done by *Reyes-Darias et al* (10). Briefly, the LBD sequence of the PctA was obtained from the *Pseudomonas aeruginosa* PAO1 genome database and the signaling region of Tar was obtained from the iGEM 2016 parts catalog (Bba K777000). These resulting constructs encode a fusion protein comprising amino acids 1 to 354 of PctA with amino acids 269 to 553 of Tar.

The LBD and HAMP of the PctA sequence together with the Tar sequence were obtained from two designed gBlocks (IDT). A double terminator sequence was obtained from the iGEM 2016 parts catalog (Bba B0015). The terminator part was digested using EcoRI and XbaI restriction enzymes resulting a backbone sequence that contains chloramphenicol resistance. The three parts were assembled on a plasmid by creating overlap regions through designed gBlocks and ligated using Gibson assembly. The chimeric chemoreceptor was transformed into Top 10 *E. coli* strain, validated and then transformed into UU1250 *E. coli* strain which lacks chemoreceptors genes *tsr, tar, tap, trg* and *aer.*

### Histamine-Tar variant design and construction

To construct the Histamine-Tar variant, a protocol presented by Moretti *et al* (14) was followed. The output of the protocol is a library of variants, ranging from dozens to thousands of protein PDB files, depending on the parameters of the design. Each variant is a mutated version of the input protein such that the mutations increase affinity to a specific ligand of our choice. By using this protocol to design a histamine sensitive Tar, a library of 870 mutated Tar receptors was obtained. Each design consisted of 3-5 iterations of the protocol for optimal results. Next, a filtering process was performed using the parameters presented in table S1 Table.

The filtering process resulted in 11 variants, which were predicted as energetically favorable to bind histamine. S1 Fig presents a sequence comparison between the designed strains and the wildtype Tar. These were constructed through mutating two places in the native Tar chemoreceptor using two sets of forward and reverse primers, as shown in S2 Table. Mutated variants were verified by sequencing and then transformed to UU1250 *E. coli* strain.

### TAR-GFP chimera construction

Tar-EGFP chimera was constructed using a flexible linker 5’-ggtagcggcagcggtagc-3’, sequence which was obtained from iGEM 2016 parts catalog (Bba J18921) to an EGFP sequence from the iGEM catalog (Bba E0040). PCR reaction to the chemoreceptors plasmid and the EGFP sequence part with an overlap region of the linker was performed, using PCR primers for EGFP sequence and chemoreceptor plasmid, as shown in S2 Table.

### Soft agar swarming assay

Swarming assay was carried out based on the Alders assay (15) (i.e. using 0.5% (w/v) soft agar plates). 5cm Petri dishes were filled with 5 mL soft agar and 5 mL of BA (bio-assay buffer) or TB based media, for PctA-Tar chimera and Histamine-Tar varianst, respectively. The plates were cooled down to room temperature in order to solidify for at least 1 h. The tested strains were grown overnight at 30°C at the appropriate media. Using a sterile tip, a drop of bacterial suspension was placed in the middle of the solidified agar containing the chemo-attractant.

### Microscope repellent chemotaxis drop assay

An overnight culture grown at 30 °C was diluted 1:50 and grown to an optical density (OD) value of ~ 1. The correlation between bacteria concentration and OD_600_ measurement was determined empirically (1 OD_600_ = 8 × 10^8^cells/mL). Cells were re-suspended in motility buffer and placed on a microscope slide (8 μL)(. A 2 μL drop of repellent was added to the bacteria a ratio of 1:5 (v/v), respectively, after validating that the final repellent concentration is not lethal for the bacteria. The bacterial response was then recorded for 15 min using Nikon Eclipse Ti microscope (magnification 100×), taking a frame every 50 ms.

### Microscope chemotaxis chip assay

An overnight culture grown at 30 °C was diluted 1:50 and grown to an OD_600_ value of ~1. A 40 μL suspension of bacteria in motility buffer was placed into a commercial sticky-Slide I Luer (Ibidi, Germany), which is specifically designed for perfusion applications. For microscopy studies, the sticky-Slides were prepared according to the manufacture’s specifications and 160 **μ**L of a chemo-effector (attractant or repellent) was added to the microfluidic channel and bacterial response was recorded for 20 min.

### Single cell fluorescence imaging

An overnight culture grown at 30 °C was diluted 1:50 and grown for 3-5 h. Soft agar gels (1% w/v) were made by dissolving 0.15 mg low melting agarose in 10 mL PBS and solidified gels were placed inside an optical plate.

10 **μ**L of the culture was then placed on the gel and incubated for 30 min in room temperature. Analysis and imaging of the cells were obtained by Nikon Eclipse Ti microscope, magnification 100×, 490 nm wavelength.

### Real-time chemotaxis on a chip

Reflectometric interference spectroscopy of two-dimensional diffraction gratings was employed to accurately measure chemotaxis. A photonic chip containing Si micron-sized wells was fixed in a custom-made flow cell heated to 37 °C and illuminated by a broadband tungsten light source (OceanOptics, LS-1) normal to the surface. Zero-order diffraction of the photonic chip was collected by a CCD spectrometer (OceanOptics, USB4000) and recorded every 30 s. All solutions were delivered to the photonic chips at a flow rate of ~0.3 mL/min using a peristaltic pump. After acquisition of a baseline in motility buffer for 45 min through the flow cell, experiments were performed in the following three stages while reflectance spectra were recorded: First, a bacterial suspension prepared in motility buffer (OD_600_ ~1) was continuously delivered until saturation of the lamellar features with bacteria. Second, the chip was washed with motility buffer solution to remove cells that were loosely adhered to the chip surface. Lastly, the chemo-effector was delivered in motility buffer for approximately 30 min. Zero-order diffraction was measured as a function of reflectance intensity versus wavelength. The frequency of the reflected light can be described using the following equation (16, 17):

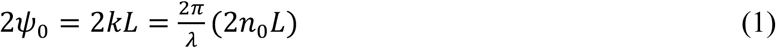

Where ψ is the phase delay between the source beam and the reflected one, λ is wavelength, *L* is the depth of the Si pores, and *n_0_* is the refractive index of the medium filling the pores. Through frequency analysis using application of a fast Fourier transform (11), the value of *2n_0_L* can be calculated. Thus *2n_0_L* refers the optical path difference, commonly termed effective optical thickness (EOT) and provides a measure for monitoring changes in refractive index that correspond to bacterial colonization within the silicon pores. Hence, upon altering the solution medium in the flow cell, a change in the refractive index occurs, leading to an increase or decrease in EOT values. (11). To normalize results, chemotaxis was measured as a function of *ΔEOT* with:

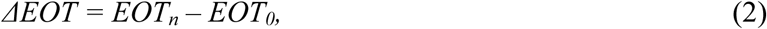

where *EOT_0_* is the *EOT* measured at *t* = 0, the time when the chemo-effector was introduced.

## Results

### Characterization of the native Tar functionality

In order to build a generic chemotactic capacity expansion toolkit, we first needed to test our rapid or digital detection approach on the natural Tar chemoreceptor. To do so, we reintroduced the *tar* gene to *E. coli* strain UU1250 that lacks all native chemoreceptors. This strain was first subjected to several chemotaxis assays, using aspartate to test the attractant response and *Ni^+^*^2^ to test the repellent response (3).

Attractant response of the Tar receptor was initially tested using the chip microscope assay (see Materials and Methods). Briefly, suspension of bacteria was placed into a commercial “sticky-Slide I Luer” Ibidi fluidic chip. The microscope was focused on a single point in the channel. Later the chemoattractant was added to the channel and the bacterial response was followed and recorded. The results (Fig. 1A-B) show that after 15 minutes the number of bacteria in the recorded frame increased from 49 to 120 after the attractant, aspartic acid, was added while the number of bacteria remained approximately unchanged (55 bacteria at time zero, and 66 after 15 minutes) for the control which was treated with motility buffer that lacks repellent or attractant. Repellent response was tested using the microscope drop assay (see Materials and Methods). Briefly, a bacterial cell suspension was placed under the microscope and a drop of repellent was added to the bacteria while recording the changes in the movement of the bacteria due to the addition. In this test, the bacteria expressing Tar immediately began to tumble in response to a repellent as expected (data not shown).

**Fig 1.**
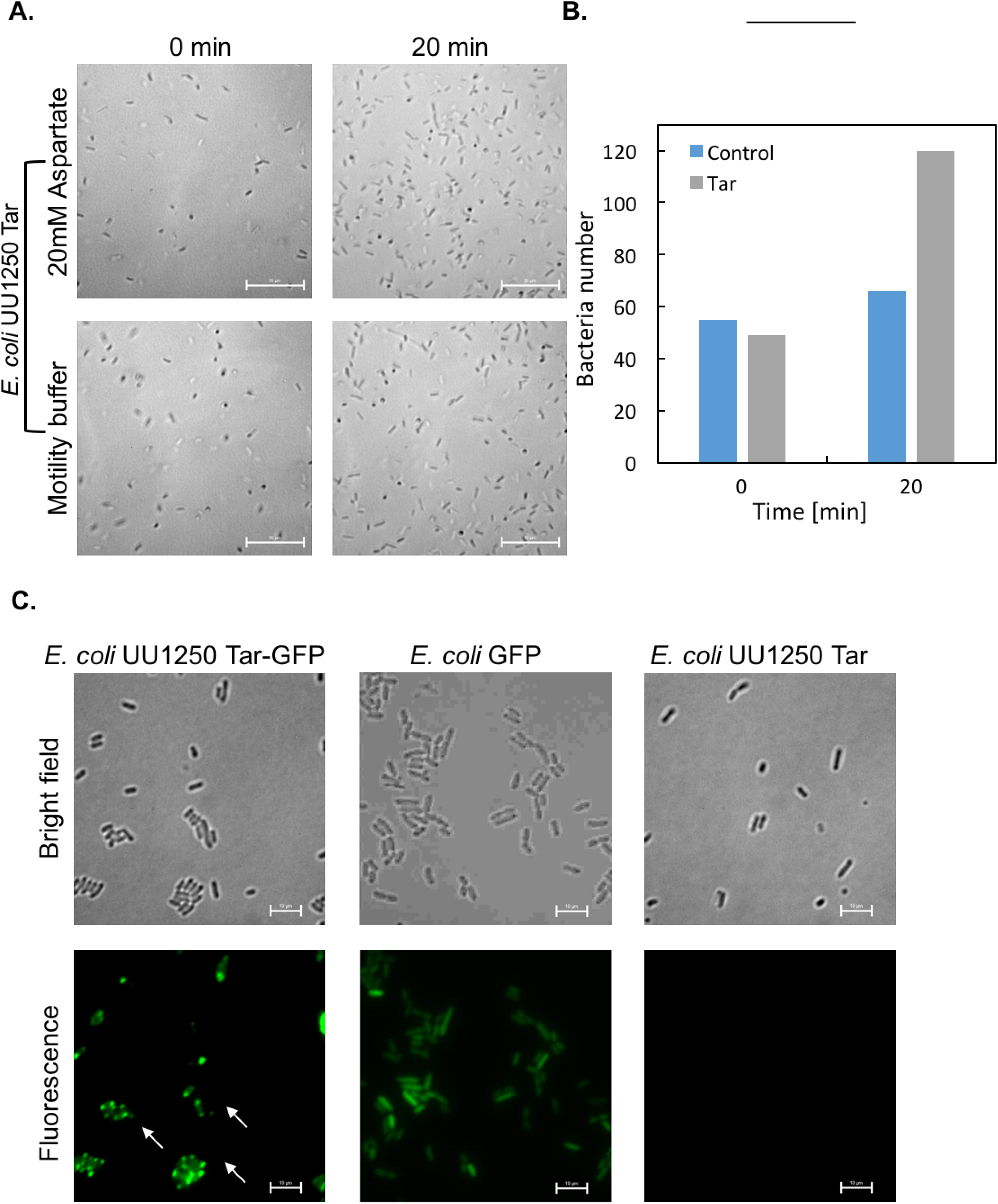
Localization of chemoreceptor at the bacteria’s membrane. (A) Optical microscope results of chemotaxis activity of the Tar variant with 10mM aspartate added, 0 and 20 minutes after adding aspartate, Control-Tar strain 0 and 20 minutes after adding motility buffer. **(B)** Quantification of the change in bacterial numbers during the chemotactic microscope attractant measurements **(C)** Bright field and GFP fluorescence (Excitation: 490 nm, Emission: 510 nm) of UU1250 strain expressing Tar-GFP chemoreceptor (upper panel) that localize in cell membrane poles, positive control-*E. coli* strain expressing GFP protein is distributed in the bacterial cell (middle panel), and negative control-UU1250 strain expressing Tar chemoreceptor (bottom panel) does not fluoresce. Y-axis corresponds to exact number of cells counted in image.

Furthermore, verification of chemoreceptor localization to the membrane poles was performed (18). This property is critical for signal amplification and adaptation of the cell, since it is crucial for additional proteins, such as kinases and adaptors to interact with the receptor, once it is situated in its proper location in the membrane (18). Although little is known about the mechanism of localization, it is important to retain this property with our newly designed receptors, to ensure a functional and sensitive chemotactic response. In Fig. 1C, we show single cell fluorescence images of bacteria expressing a chimera of the Tar-GFP-chemoreceptor (18, 19). The figure shows that Tar-GFP (upper panel) can be detected in the cell poles, indicating a proper migration and localization of the Tar receptor whereas the strain expressing GFP only lacks GFP translocation to the bacterial cell membrane (middle panel) and the strains expressing Tar only (bottom panel) did not display any GFP fluorescence.

### Real-time chemotaxis-on-a-chip

Though numerous methods have been developed to study chemotactic responses, none provide a real-time measurement without the use of fluorescence labeling. In recent years, the use of porous Si matrices for bio-sensing applications has readily increased and has encompassed the detection of various analytes such as DNA, proteins, and bacteria (20-22). Thus, we decided to employ the specific “trap and track” method of bacteria within Si diffraction gratings demonstrated by Massad-Ivanir et al. (11, 23) as a proof-of-concept basis to monitor chemotactic responses in real-time using three different bacterial strains (see Table 1). Figure 2A schematically illustrates the concept of the chemotaxis-on-a-chip assay, where the optical response of photonic chip, comprised of a Si diffraction grating with square wells ~3.7 μm in width (Fig. 2B), is monitored.

**Fig 2.**
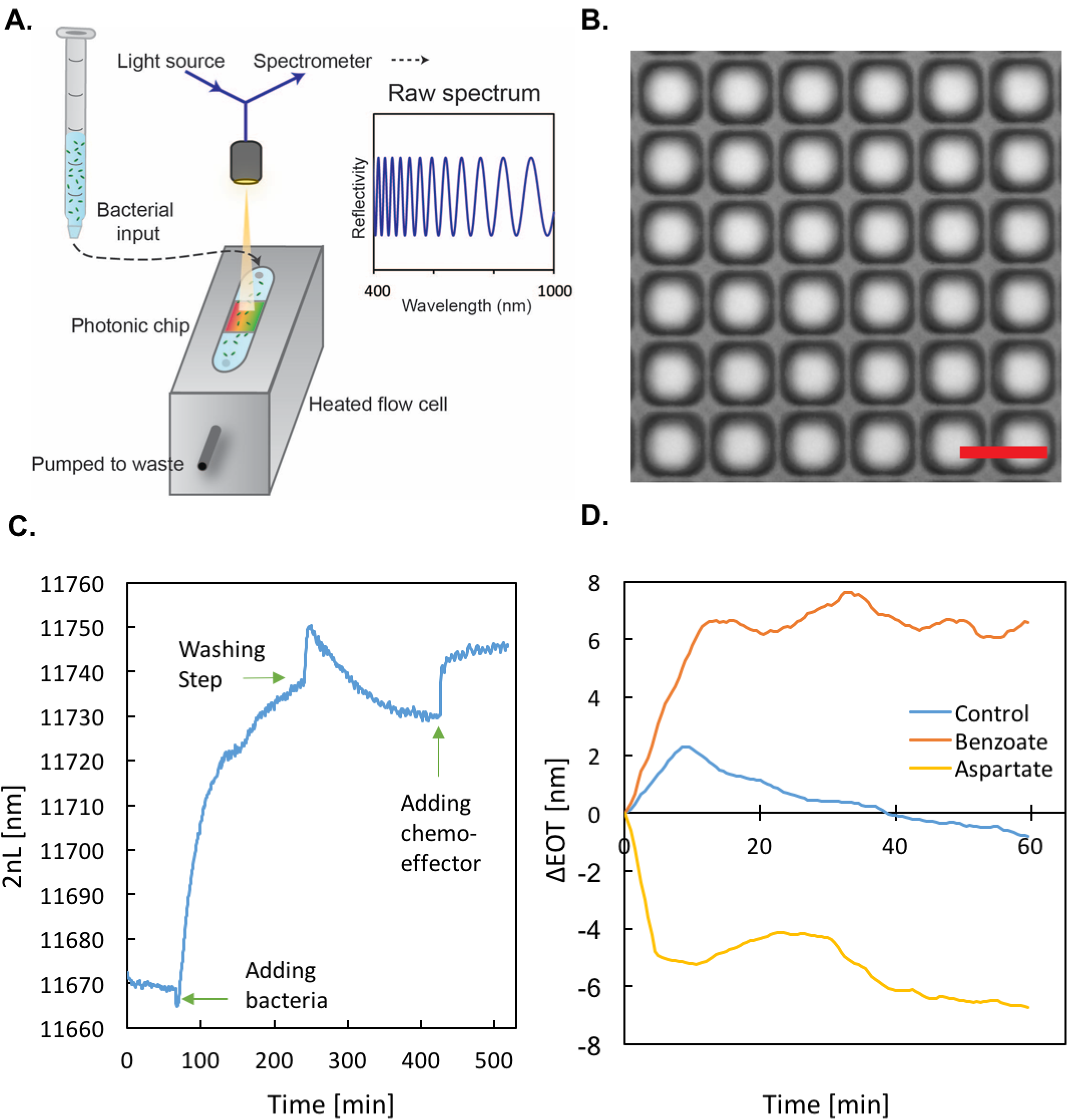
Chemotaxis-on-a-chip concept and result. (A) Schematic of the real-time chemotaxis-on-a-chip system in which a photonic chip, housed in a heated flow cell is filled with various suspensions while the zero-order diffractive reflectance of the chip is monitored. **(B)** Optical microscope image of the photonic chip, which is comprised of a Si diffraction grating. Scale bar represents 5 μm. **(C)** Optical read-out for a single experiment, presenting all three experimental steps: (i) Adding bacterial solution. (ii) Washing with motility buffer. (iii) Adding a chemo-effector. **(D)** Chemotaxis monitoring presenting three chemotactic responses: ΔZras with repellent containing 50 mM sodium benzoate (orange), UU1250 control with repellent containing 50 mM sodium benzoate (blue), native Tar with attractant containing 20 mM aspartate (yellow).

Results for a typical chemotaxis assay are presented in Figure 2C in terms of the recorded EOT value vs. time. The panel represents the full EOT recorded for the duration of an entire experiment (starting from baseline acquisition) over time for the bacterial strain with as the chemo-effector. After the addition of the bacteria, an EOT shift of ~70 nm occurred over the span of 200 minutes. Subsequent washing of the chip with motility buffer led to a decrease in EOT, as excess, non-adhered cells are removed. Introduction of the chemo-effector (50 mM benzoate) results in a sharp increase in the EOT value, which is ascribed to higher refractive index of the solution (e.g., when chemoattractant was added). Notably, while the preparation of the chip to chemotaxis experiments is time consuming (~ 7 hours), a positive chemotactic response is recorded within approximately 10 minutes.

To observe differences in chemotactic responses, three different bacterial strains were tested (Table 1), and ΔEOT measurements are plotted in Figure 2D. First, the response of the bacterial strain expressing the Tar chemoreceptor was studied upon the introduction of its natural attractant, aspartate (20 mM in motility buffer). In this case, a rapid decrease in the EOT (of ~4 nm) is observed within less than 5 min, after which a continuous moderate reduction is recorded (Fig. 2D, yellow trace). This behavior is ascribed to the decreasing population of cells within the chip, as the bacteria respond to the chemoattractant gradient in the solution and actively move towards it (6). An opposite trend is obtained when the Δ*Zras* strain, which expresses all four natural chemoreceptors, is exposed to the chemo-repellent solution of benzoate (50 mM in motility buffer). A rapid increase in the EOT is observed within 5 min, followed by a stabilization of the EOT signal around a constant value within the next hour (Fig. 2D, orange trace). The response of the UU1250 strain, which lacks all chemoreceptors, to benzoate was also tested as control. Because this strain lacks the ability to move towards the chemo-effector, most bacteria remained inside the pores, resulting in small amplitude fluctuations of ΔEOT around the baseline EOT value (Fig. 2D, blue trace). Thus, the Tar and ΔZras strains, which are chemotactically active, behaved as expected, either leaving the pores or entering them depending on the effector introduced, resulting in a drop or rise in EOT, respectively.

**Table 1.** Bacterial strains and chemo-effectors tested using the chemotaxis-on-a-chip platform.

| Bacterial strain | Chemoreceptors | Chemo-effector tested |
| --- | --- | --- |
| *E. coli Tar* | Tar chemoreceptor | 20 mM aspartate (attractant) |
| *E. coli ΔZras* | 4 chemoreceptors | 50 mM benzoate (repellent) |
| *E. coli UU1250* | none | 50 mM benzoate (repellent) |

### PctA-Tar chimera exhibits chemotactic responses towards PctA specific chemo-effectors

As a first attempt to test synthetic chemo-receptors, we constructed a chimeric chemoreceptor from *P. Aeruginosa* PctA chemoreceptor and *E.coli’s* Tar. A different variant of this chimera was previously demonstrated to be chemotactic in *E. coli* by Reyes-Darias et al (10). PctA mediates chemotactic movement towards all L-amino acids (except aspartic acid) and away from organic compounds such as TCE (10). To generate the chimeric receptor PctA-Tar we chose to use the ligandbinding as well Histidine kinase, adenylyl cyclase, methyl-accepting -taxis proteins and phosphatase (HAMP) domains of PctA and fused it to the Tar signaling domain. The resulting plasmid was transformed into the chemotactic null *E. coli* UU1250 strain.

To confirm the functionality of the chimeric receptor, a swarming plate assay was carried out. In this assay, a drop of a bacterial suspension is placed in the middle of a soft agar plate containing the chemo-attractant. Subsequently, bacteria metabolize the nutrients in their immediate surroundings thus a concentration gradient of the chemo-attractant is formed causing the bacteria to move through the pores of the agar, and advance towards higher concentrations of nutrients. As the bacteria progress through the gel, “chemotactic rings” are formed and can be observed after overnight incubation at 30 °C. In Fig. 3, no “chemotactic ring” formation can be observed for the control strain UU1250 (Fig. 3B, center) indicating the lack of chemotactic movement. In contrast, “chemotactic ring” formation can be seen for the chimeric PctA-Tar receptor (Fig. 3B, left) as well as for the enhanced motility chemotactic strain *ΔZras* (Fig. 3B, right), confirming the recovery of chemotactic ability for the chimeric PctA-Tar receptor in the UU1250 strain of *E. coli.* To show that the PctA-Tar chimera has the ability to translocate to the membrane poles, GFP was fused to its C-terminus and microscope images of GFP localization were recorded. S2 Fig confirms the polar localization of the PctA-Tar chimera as the constructed chemoreceptor is localized to the membrane poles.

**Fig 3.**
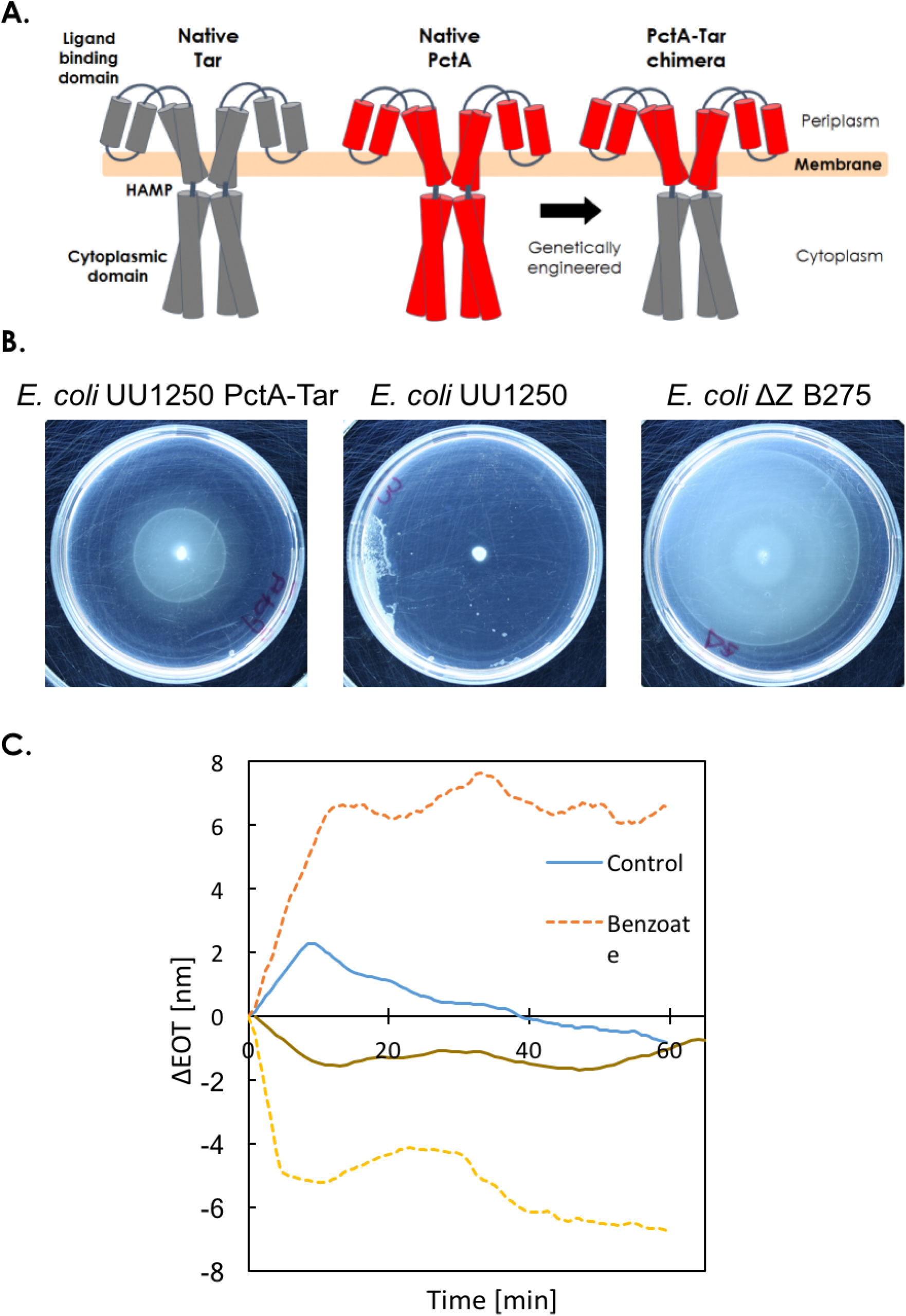
PctA-Tar chimeric receptor construction and functional analysis. **(A)**Schematic representation of the native Tar chemoreceptor (left), native PctA of *Pseudomonas aeruginosa* receptor (middle) and PctA-Tar chimera (right) that contains the PctA ligand binding domain and the Tar signaling domain. Scheme has been adapted from Reyes-Darias *et al* (10). **(B)** Swarming assay was performed for the attractant response of the PctA-Tar chimera, the negative control strain UU1250 and the positive control strain ΔZras. Plates were incubated at 30 °C for 24 hr. (C) Real time chemotaxis measurement results presenting four chemotactic responses: ΔZras with repellent containing 50 mM sodium benzoate (orange), PctA-Tar chimera with attractant containing 10mM alanine (brown), UU1250 control with repellent containing 50 mM sodium benzoate (blue), native Tar with attractant containing 20 mM aspartate (yellow). See also S3 Fig.

To further characterize the PctA-Tar synthetic chemoreceptor, we tested its response to the L-amino acid Alanine attractant using our real time chemotaxis assay (Fig. 3C). The graph presents the ΔEOT measurements obtained for the PctA-Tar strain when subjected to an attractant gradient of 10 mM Alanine (brown). The Alanine ΔEOT trace is plotted together with the traces obtained for the Tar control shown in Fig. 2D. The results show that while a rapid (~10 minutes) and consistent reduction in ΔEOT is observed for Alanine, consistent with it being an attractant, the overall amplitude of the response is ~x5 weaker than what was previously observed for Tar. However, the dose response curve generated by the PctA-Tar strain for alanine, characterized by the speed of the response (~5-10 minutes) and the stabilization around a lower value of ΔEOT is consistent with dose response obtained for aspartate with the enhanced motility *ΔZras* strain. As a result, the discrepancies in the dose response curves can be explained with by either the enhanced motility of *ΔZras,* or by a lower sensitivity to the Alanine gradient by the PctA-Tar chimera.

### Computationally designed histamine-sensitive chemoreceptor shows attraction response to histamine

The bacterial world offers a relatively small selection of chemoreceptors in comparison to the vast number of possible ligands. This fact alone limits the capacity to generate synthetic chemoreceptors using a chimeric strategy as shown above. To overcome these limitations and expand the chemotaxis repertoire of *E. coli* beyond known naturally occurring ligand binding domains, we attempted to apply a computational design approach to construct novel chemoreceptors. We chose, as a proof of concept, to construct a histamine binding receptor, as it is a small molecule and a variant of an amino acid, histidine. Histamine is known to be found in decaying food, especially rotten fish, and can cause a food-allergy-like immune response(24), which in some cases can be life threatening. To construct a histamine-sensitive receptor, we used the Rosetta software suite, a collection of algorithms and programs for macromolecular modeling and design. For the generation of new ligand-binding domains we followed the protocol described by Moretti *et al* (14). Briefly, the program is given two inputs, A PDB file containing the sequence of the ligand binding domain of the receptor, and the chemical structure of the target ligand. The program then inserts random mutations into the sequence of the native binding domain and calculates the effects of these mutations in terms of energy and binding affinity. The output of a single protocol iteration is a library of dozens to hundreds of variants, each with its own set of mutations. For each target ligand, we ran 3-5 iterations of the protocol, to increase the chance of a successful hit. Subsequently, the virtual library was subjected to a filtering process designed to eliminate variants which Rosetta predicts will not bind the ligand correctly, this is done according to the parameters which are specific per design (see Table S1). Using this algorithm, we managed to generate a library of mutated Tar ligand binding domains that were computational predicted to bind histamine and activate the chemotaxis pathway in response to it. Out of the 870 suggested mutations, only 11 variants passed the filtering process, meaning, they were predicted to bind histamine. As expected, all mutations are located near the natural binding pocket for aspartate as can be seen in Fig. 4A and in Fig S1. Each color in the three-dimensional plot of the Tar ligand binding domain represents the mutations of each variant, presented in their unbound form.

**Fig 4.**
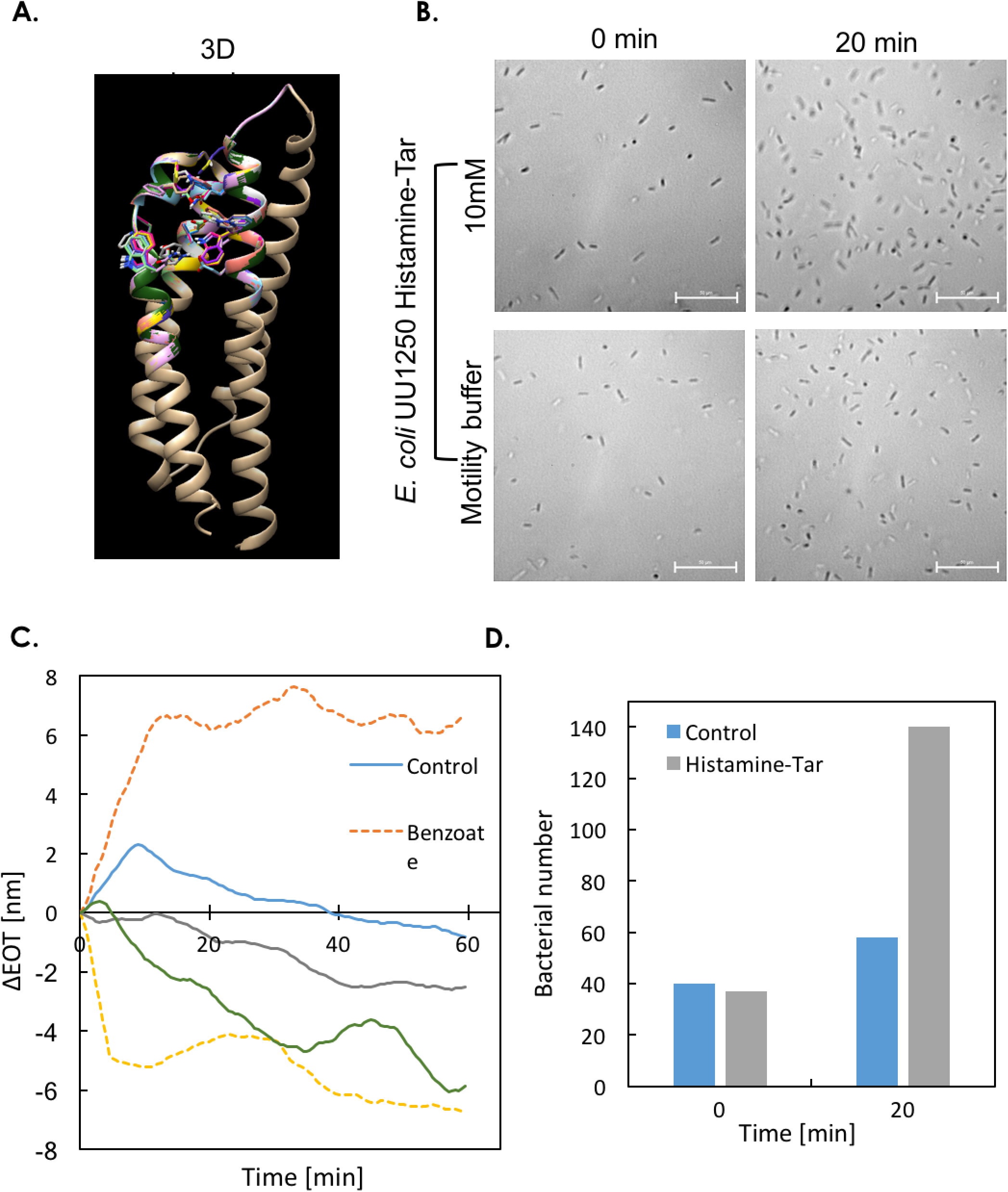
Histamine-Tar variant construction and functional analysis. (A) 3D imaging of the 11 mutated variants together with the native Tar (wild type) located near the binding pocket, each color represents a different variant **(B)** Microscope results of chemotaxis activity of the histamine-Tar variant with 10mM histamine added, 0 and 20 minutes after adding histamine, Control-histamine-Tar strain 0 and 20 minutes after adding motility buffer. Lower graph showsquantification of the change in bacterial numbers during the chemotactic microscope measurements. **(C)** Chemotaxis monitoring presenting five chemotactic responses: AZras with repellent containing 50 mM sodium benzoate (orange), UU1250 control with repellent containing 50 mM sodium benzoate (blue), control of histamine-Tar variant with attractant containing 20 mM aspartate (grey), histamine-Tar variant with attractant containing 10 mM histamine (green), native Tar with attractant containing 20 mM aspartate (yellow).

Next, these 11 variants were cloned into the native Tar ligand-binding domain and tested for chemotactic activity towards histamine, using a bacterial suspension in motility buffer was placed into a commercial Ibidi fluidic chip. Histamine was added and the bacterial response was recorded. Out of the 11 tested variants (data not shown), one variant (histamine-Tar#9) mediated an attraction response towards histamine. 20 minutes after addition of histamine, the bacterial concentration increased more than two-fold (Fig. 4B) close to where the histamine has been added to the slide, whereas the same strain did not move towards motility buffer thus demonstrating the first example of a newly generated chemoreceptor using computational design. To show that the histamine-Tar variant retains the ability to translocate to the membrane poles, GFP was fused to its C-terminus and microscope images of GFP localization were recorded. S2 Fig confirms the polar localization of the Histamine-Tar variant as the constructed chemoreceptor is localized to the membrane poles.

Finally, we tested the histamine-Tar variant using our real time quantitative chemotaxis assay. The results are plotted in (Fig. 4C). The graph presents the dose response as function of time of histamine-Tar to 20 mM aspartate as a control (grey) and 10 mM histamine as attractant (green). The figure shows that the mutated variant displays a delayed “attractant” response to aspartate, which possibly reaches steady-state ΔEOT levels after 45 minutes, but with a final amplitude that is ~3x smaller than the natural Tar’s response to Aspartate. The histamine-Tar response to addition of histamine results in a moderate and continual change of ΔEOT ultimately reaching an amplitude similar to what was observed for the natural Tar’s response to aspartate with steady state possibly being reached at 30 m. Consequently, while the dose response of the mutated Tar to histamine is observed, the speed of the response is 4-5x slower than Tar’s response to asparate indicating that additional mutations may be needed in order to improve the chemotactic behavior of this mutant.

## Discussion

The bacterial chemotaxis system has been studied for several decades due to its wide range of applications. In recent years, with the advance of technology, knowledge of the inner working of this system has led to attempts to harness it for targeted usage via different approaches, such as: rational design of chimeras and directed evolution (10, 8). In this work, we showed using a combined computational and a real-time quantitative approach that the *E. coli* Tar chemoreceptor can be respecified to a new ligand that does not have a known chemoreceptor in the microbial metagenome.

A reliable computational design algorithm can complement synthetic chimera strategies (as demonstrated by Reyes-Darias *et al* (10) and repeated by us here), in two ways: first, construction of a chemoreceptor chimera can be sensitive to domain compatibility with other receptors, mainly due to lack of knowledge regarding other bacterial chemoreceptors. While a recent paper by the same group (7) portends to have developed a generic approach for assembly of chimeras, it only has been demonstrated on a handful of new chimeras, and is thus not completely full proof. Second, computational design algorithms such as the Rosetta software are not limited to the natural milieu of chemoreceptors, and thus can be used to generate completely new receptors as was shown here.

In order to utilize this approach and to systematically mine the potential ligand space for new nonnatural receptors, we developed a new quantitative, real-time, and digital-like chemotaxis detection assay. Our approach is an adaptation of the “trap and track” method of bacteria within Si diffraction gratings demonstrated by Massad-Ivanir et al. (11, 23). Here, using a steady flow, we generated ligand concentration gradient inside individual porous Si wells, which force the bacteria to either leave the well in case of an attractant or congregate at the bottom if the ligand is a repellant. Detection of motion is carried out via the change in refraction index or EOT, which facilitates a rapid and quantitative characterization of the chemotactic response. In this work, we studied three different chemoreceptors using three separate attractants and one repellant. Our results shows that each ligand/chemoreceptor pair can be characterized based on the amplitude of refractive index change, the length of time that is required for steady-state level to be reached, and a digital classification into attractant/repellant depending on whether the change in refractive index is negative or positive respectively. Consequently, our quantitative, real-time, and digital approach provides additional read-out into the ligand-chemoreceptor interaction that can facilitate better characterization of chemotaxis. Finally, we envision that are method can potentially be used to rapidly characterize the response of synthetic chemoreceptors to novel ligands, thus facilitating a methodology where dozens of new ligands and chemoreceptor pairs can be scanned, characterized, and even improved with complementary directed evolution-based approaches. Such a tool will not only contribute to the advancing of our understanding of chemotaxis, but also provide the means by which to generate completely synthetic chemotactic bacteria for a whole range of applications from bio-remediation to medical diagnostics.

## Acknowledgments

We would like to thank the Department of Biotechnology and Food Engineering at the Technion for allocating laboratory space. We gratefully acknowledge the financial support of the Lorry I. Lokey Interdisciplinary Center for Life Sciences and Engineering, Prof. Peretz Lavie - president of the Technion, and Israeli Ministry of Science, Technology and Space for funding. We are also thankful to Dr. Nadav Ben Dov and Dr. Naama Massad-Ivanir from the Department of Biotechnology and Food Engineering for their guidance and advice.

## Supporting information

**S1 Fig.**
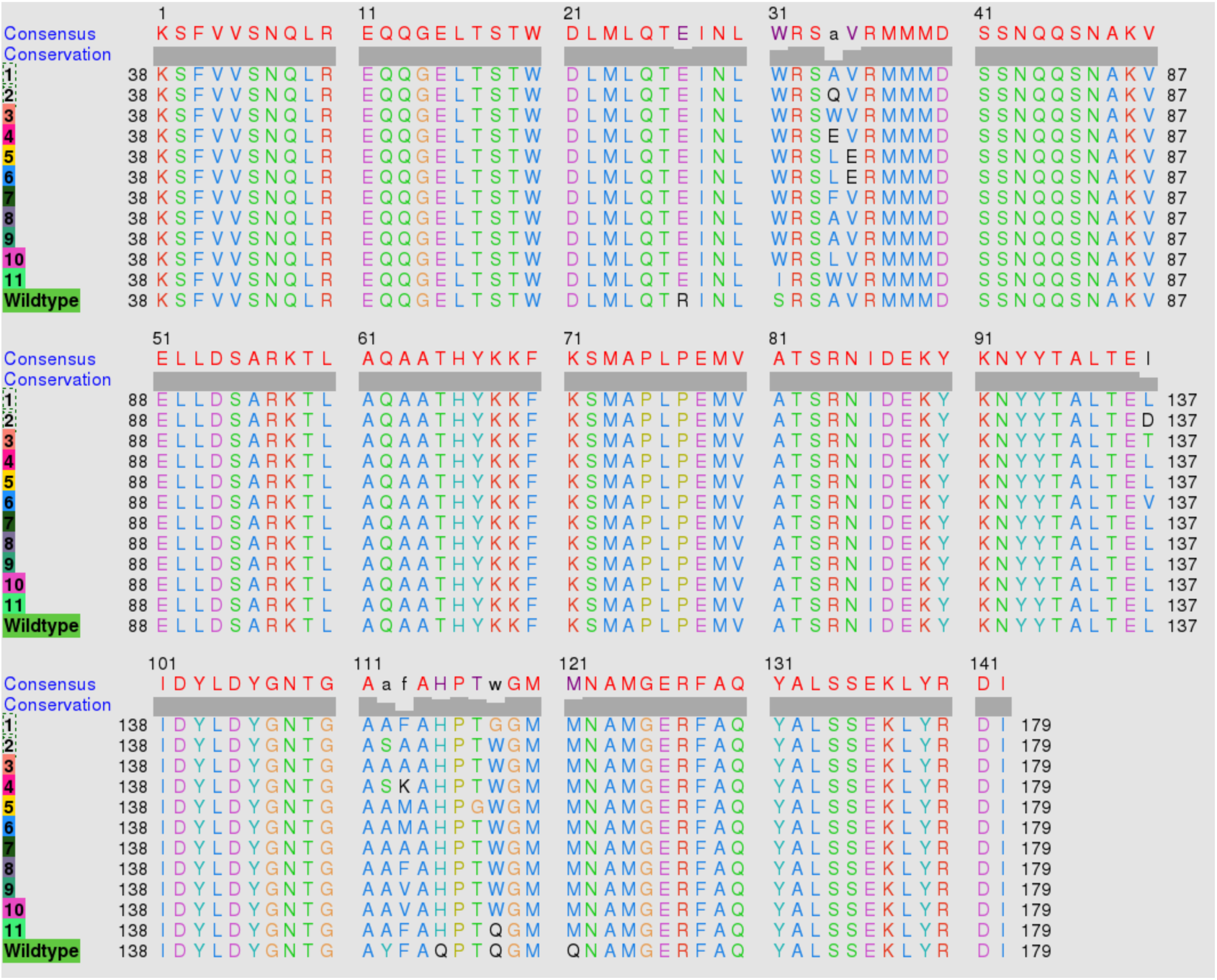
Amino acid sequences of the ligand binding domains. Amino acid sequences of the ligand binding domains of the 11 mutated variants which passed the filtering process, together with that of the native Tar (wild type).

**S2 Fig.**
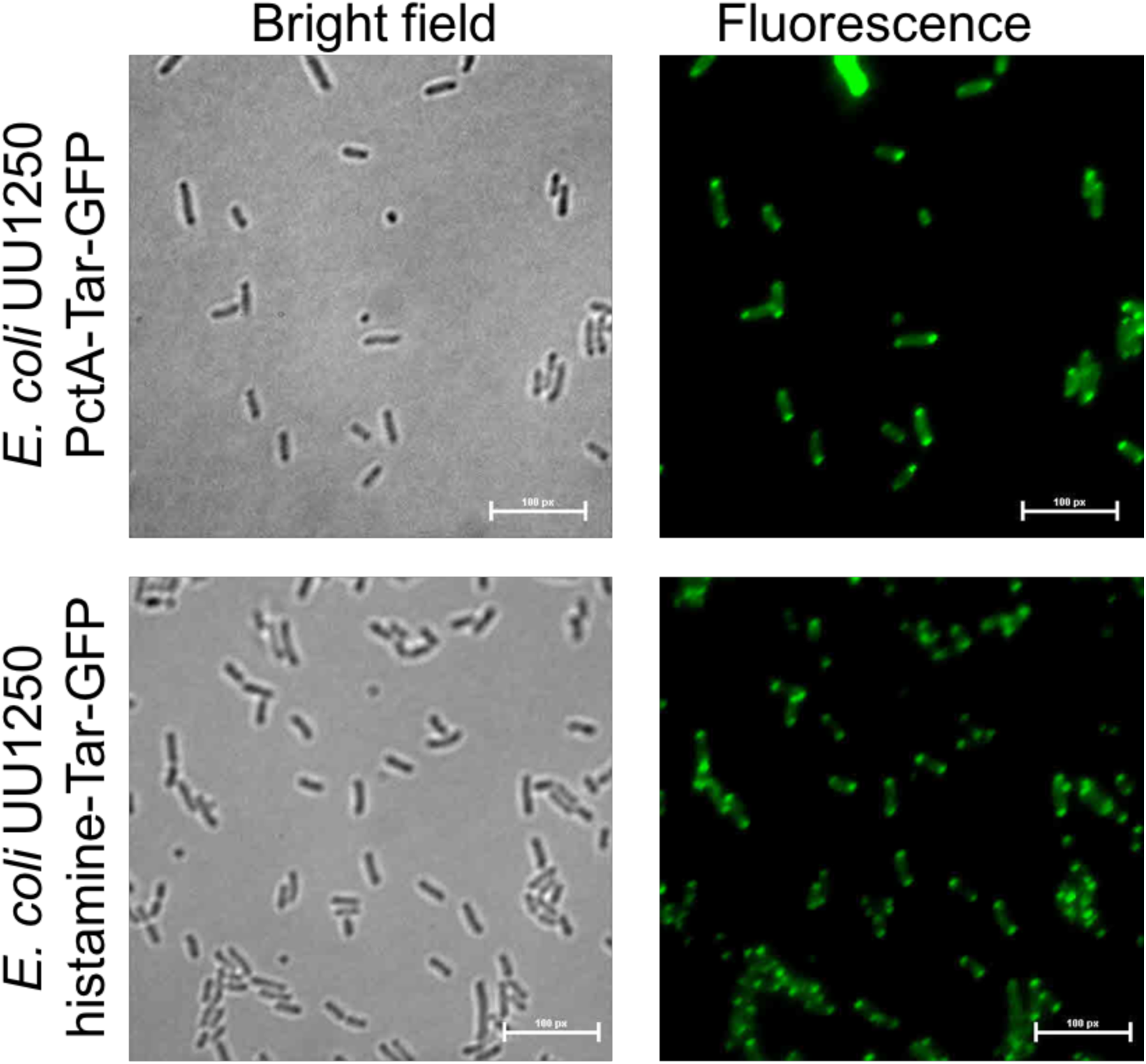
Bright field and GFP fluorescence. (Excitation: 490 nm, Emission: 510 nm) of UU1250 strain expressing PctA-Tar-GFP chemoreceptor (upper panel) localize in cell membrane poles, and UU1250 strain expressing PctA-Tar-GFP chemoreceptor (bottom panel) localize in cell membrane poles.

**S3 Fig.**
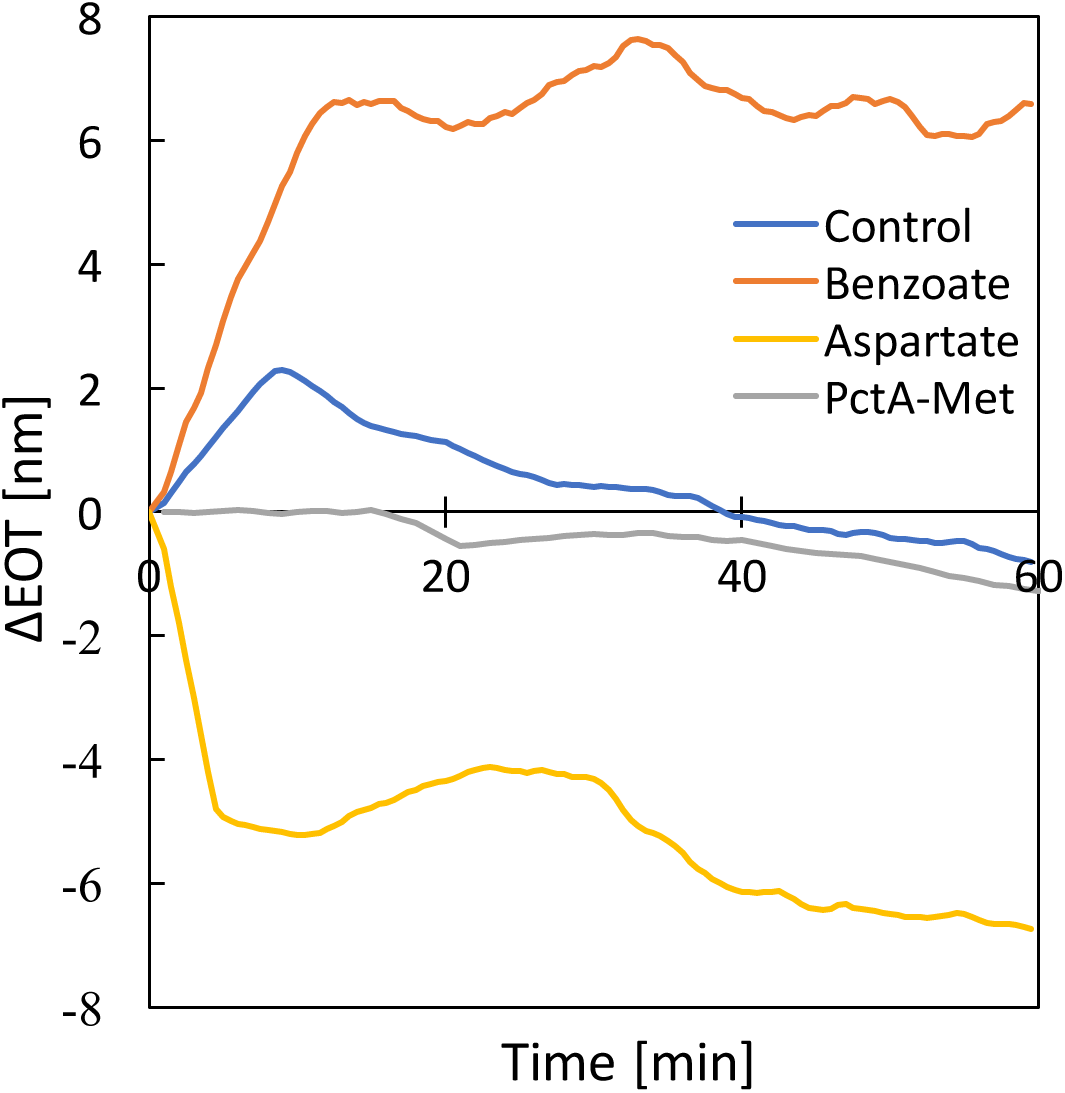
Real time chemotaxis measurement results presenting four chemotactic responses: AZras with repellent containing 50 mM sodium benzoate (orange), PctA-Tar chimera with attractant containing 10mM methionine (brown), UU1250 control with repellent containing 50 mM sodium benzoate (blue), native Tar with attractant containing 20 mM aspartate (yellow).

**S1 Table.** Filtering parameters. Filtering parameters used according to Rosetta protocol.

| **Name of Parameter** | **Meaning** |
| --- | --- |
| **total_score** | weighted average of all parameters |
| **Transform_accept_ratio** | How well the low-resolution Monte Carlo stage worked. It is a number between 0 and 1 and is provided as a diagnostic |
| **fa_elec** | Coulombic electrostatic potential |
| **fa_intra_rep** | The repulsive energy within residues |
| **fa_pair** | Statistical residue-residue interaction potential. Very useful in protein-protein interactions |
| **fa_rep** | The repulsive portion of the Lenard-Jones potential (11). If high, then there are clashes in the protein |
| **if_X_hbond_bb_sc**<br>**if_X_hbond_sc** | Hydrogen bonding term between protein backbones and side chain (ligand) |
| **interface_delta_X** | The energy of interactions between the ligand (chain X) and the protein |
| **ligand_is_touching_X** | 1 if ligand is close to the protein, 0 otherwise |
| **complex_normalized** | Total score of the protein complex with the ligand (holo state) normalized by the number of residues |
| **dG_cross** | The energy of interaction between the two sides of the complex, calculated in the holo state (protein bound to ligand) |
| **dG_separated** | The energy of interaction between the two sides of the complex, calculated by taking the score of the holo state, then separating the two sides of the interface, optionally repacking, and then calculating the score in the separated state. This is particularly useful if you think you have an "induced fit" type situation, and want to correct for repacking in the absence of the ligand. |
| **delta_unsatHbonds** | How many unsatisfied hydrogen bonds are introduced by the design. Unsatisfied hydrogen bonds are ones where the atom is able to make a hydrogen bond but does not because it's blocked by other residues |
| **Packstat** | How well the protein is packed. values between 0-1 with 1 being better |
| **sc_value** | Shape complementary score. Values between 0-1 with 1 being better. It looks at the correspondence between normal vectors of the surfaces of the protein and ligand |
| **side1_normalized**<br>**side2_normalized** | The energies of the two sides of the interface, possibly normalized by the number of residues |

**S2 Table.**
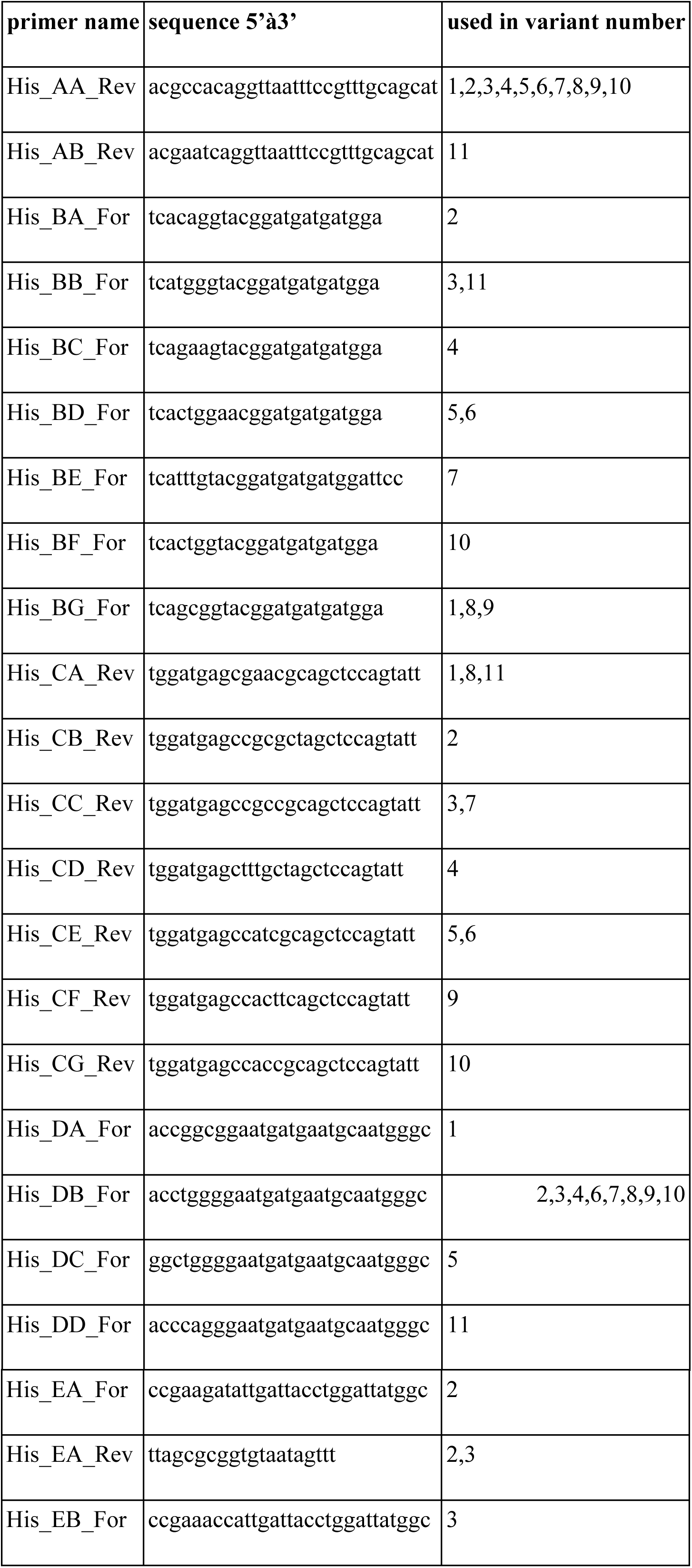
List primers used to construct Histamine-Tar variants and Histamine-Tar-GFP chimera.

